# FamPlex: a resource for entity recognition and relationship resolution of human protein families and complexes in biomedical text mining

**DOI:** 10.1101/225698

**Authors:** John A Bachman, Benjamin M Gyori, Peter K Sorger

**Affiliations:** Laboratory of Systems Pharmacology, Harvard Medical School, 200 Longwood Ave, 02115 Boston, MA, USA

**Keywords:** Text mining, protein families, grounding, named entity linking, named entity recognition, biocuration, event extraction, natural language processing

## Abstract

**Background:** For automated reading of scientific publications to extract useful information about molecular mechanisms it is critical that genes, proteins and other entities be correctly associated with uniform identifiers, a process known as named entity linking or “grounding.” Correct grounding is essential for resolving relationships among mined information, curated interaction databases, and biological datasets. The accuracy of this process is largely dependent on the availability of machine-readable resources associating synonyms and abbreviations commonly found in biomedical literature with uniform identifiers.

**Results:** In a task involving automated reading of ∼215,000 articles using the REACH event extraction software we found that grounding was disproportionately inaccurate for multi-protein families (e.g., “AKT”) and complexes with multiple subunits (e.g.”NF-*κ*B”). To address this problem we constructed FamPlex, a manually curated resource defining protein families and complexes as they are commonly encountered in biomedical text. In FamPlex the gene-level constituents of families and complexes are defined in a flexible format allowing for multi-level, hierarchical membership. To create FamPlex, text strings corresponding to entities were identified empirically from literature and linked manually to uniform identifiers; these identifiers were also mapped to equivalent entries in multiple related databases. FamPlex also includes curated prefix and suffix patterns that improve named entity recognition and event extraction. Evaluation of REACH extractions on a test corpus of ∼54,000 articles showed that FamPlex significantly increased grounding accuracy for families and complexes (from 15% to 71%). The hierarchical organization of entities in FamPlex also made it possible to integrate otherwise unconnected mechanistic information across families, subfamilies, and individual proteins. Applications of FamPlex to the TRIPS/DRUM reading system and the Biocreative VI Bioentity Normalization Task dataset demonstrated the utility of FamPlex in other settings.

**Conclusion:** FamPlex is an effective resource for improving named entity recognition, grounding, and relationship resolution in automated reading of biomedical text. The content in FamPlex is available in both tabular and Open Biomedical Ontology formats at https://github.com/sorgerlab/famplex under the Creative Commons CC0 license and has been integrated into the TRIPS/DRUM and REACH reading systems.

## Background

A critical challenge in contemporary molecular biology is integrating detailed mechanistic information about specific genes and proteins with genome-scale information about the interaction networks in which these genes participate. Networks of molecular mechanisms are powerful tools for interpreting large-scale data in the context of prior knowledge [1, 2, 3, 4]. The construction of biological networks benefits from exchange formats such as BioPAX [5] that allow disparate databases to be aggregated into uniform, machine-readable resources such as Pathway Commons [6]. However, a significant fraction of the information available in the literature has not been recorded in pathway databases. Text mining has the potential to address this gap by augmenting curated network resources with molecular mechanisms automatically extracted from the literature. However, current systems are not yet able to extract mechanisms with a quality equal to that of human curators [7].

One challenge in using text-mined information for biological data analysis is that molecular mechanisms are often described in the literature in terms of aggregate entities such as multi-protein families (e.g., “RAS”, “AKT”) and multi-subunit complexes (e.g., “NF-*κ*B, “AP-1”) rather than the specific genes or proteins measured in large-scale experiments. For example, a Pubmed search for “NF-kappaB” yields over 64,000 citations; this transcription factor is not a single molecular entity but rather a class of heterodimers involving combinations of at least five different genes in two families (*RELA, RELB, REL, NFKB1*, and *NFKB2*). This poses two challenges for machine reading. First, the text string “NF-*κ*B” must be normalized to a standard identifier, a task known variously as named entity linking (NEL), named entity normalization (NEN), named entity disambiguation (NED), or simply “grounding.” [8]. Second, the mapping of “NF-*κ*B” to its constituents must be established so that the activities of NF-*κ*B can be linked to the properties of the genes from which it is comprised. Such “static relations” must be resolved either by explicit curation or algorithmically [9, 10, 11].

Success in the first task, grounding, is essential for practical applications of text mining [12, 13]. Entities without associated identifiers cannot be used for down-stream assembly and interpretation tasks, and systematic *misidentification* of entities clutters extracted networks with errors that skew data analysis. Relevant approaches to grounding have been studied extensively in the context of the general problem of biomedical entity normalization [14, 8, 15, 16], and generally involve two steps. First, a named entity as encountered in text is normalized, for example by stemming [17], removal of affixes [10], or expansion of abbreviations [16]. Effective preprocessing depends on an explicit or implicit representation of how specific entities (e.g., diseases vs. chemicals vs. genes) variously appear in text (see 2.2.4 in [16]).

The normalized string is then matched to names and synonyms in existing taxonomies [13]. Difficulties in grounding protein families and complexes are encountered in this latter step because there is no standard ontology for these entities as they are commonly described in the scientific literature. Relevant identifiers can be found in protein family databases (InterPro, PFAM, NextProt) and curated interaction databases (Reactome, SIGNOR, OpenBEL) allowing complexes and families to be resolved into their constituent genes. However, such databases generally lack lexical synonyms corresponding to the many ways in which entities are referenced in text, limiting their value for literature mining. Conversely, general biomedical vocabularies and thesauri such as NCIT and MeSH contain entries and lexical synonyms for families and complexes but often lack the ontological resolution of these terms into child concepts (e.g. entries C94701 in NCIT and D055372 in MeSH for the holo-enzyme AMPK, neither of which define its constituents). In combination, these diverse databases provide substantial information about families and complexes, but integration of this information is difficult because they rarely contain cross-references for related concepts among themselves. Prior work has addressed aspects of normalization for protein families, for example by automatically identifying families and their constituents directly from the literature [9, 15] or by combining information in gene family databases with patterns in the names and synonyms of genes [10, 18]. However, the problem of identifying, normalizing, and linking information about protein families and complexes is less well-understood than that of gene normalization [8, 18, 16], and draws on a smaller base of taxonomic resources. In this paper we describe FamPlex, a curated lexical and ontological resource that improves grounding and relationship resolution for families and complexes encountered in the mining and curation of biomedical text. FamPlex contains a set of identifiers for protein families and complexes along with mappings that link: (i) text strings and FamPlex identifiers, (ii) FamPlex identifiers and identifiers representing protein families and complexes in other resources, and (iii) FamPlex families/complexes and their constituent members. FamPlex also contains a list of prefixes and suffixes frequently appended to protein names for use in named entity recognition (NER) and entity normalization. The FamPlex resource consists of a set of comma-separated value (CSV) files listing entities and relations, along with Python scripts for checking consistency and identifying equivalent identifiers in other databases. FamPlex is hosted on GitHub at https://github.com/sorgerlab/famplex and is made available under the Creative Commons CC0 license. It is also available in the Open Biomedical Ontology (OBO) format and can be accessed via the NCBO BioPortal [19] at http://bioportal.bioontology.org/ontologies/FPLX.

## Construction and Content

Development of FamPlex was motivated by an empirical analysis of grounding accuracy in events extracted by the REACH biomedical literature mining software [20, 21]. As described in detail below, we found that grounding of protein families was disproportionately inaccurate and that a relatively small proportion of frequently misgrounded entities accounted for the bulk of all grounding errors. An examination of existing resources highlighted the fragmented nature of information on protein families and complexes and the general lack of suitability of these resources for literature mining. FamPlex was conceived as a a “bridging” resource to link available information about families, complexes, and other frequently misgrounded entities across a diverse set of existing bioinformatics databases.

At the core of FamPlex is a set of identifiers representing protein families and complexes (Figure 1A). FamPlex represents the hierarchical relationships of these high-level entities to each other and to individual genes, along with corresponding synonyms in text and cross-references to other databases where available. Entities and mappings are recorded in a set of CSV files.

**Figure 1.**
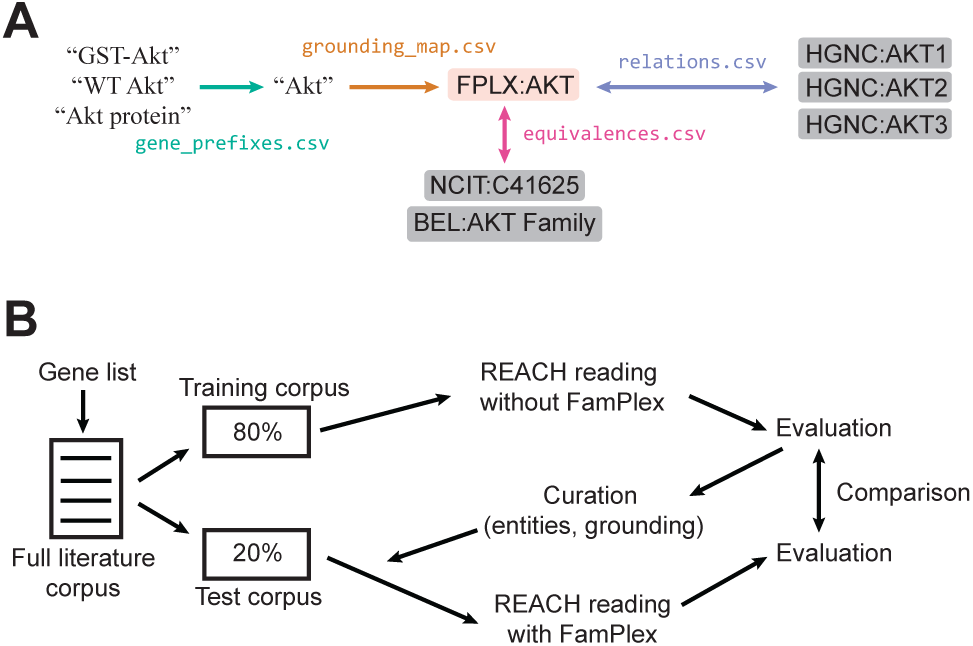
FamPlex links named entities to protein families and complexes and their constituents. **(A)** Structure of FamPlex content. The affixes in `gene prefixes.csv` can be used to improve recognition of molecular entity names, which can be linked to database identifiers using the lexical synonyms in `grounding map.csv`. FamPlex itself contains identifiers representing families and complexes which are mapped to corresponding identifiers in other databases in `equivalences.csv`. Hierarchical relationships among families, complexes, and genes are listed in `relations.csv`. **(B)** Workflow for curation and evaluation. A gene list was used to define a corpus of articles that was divided into two subsets, “training” and “test”. The “training” corpus was processed with REACH and results were evaluated and used to guide curation. The “test” corpus was processed after incorporation of FamPlex and results were compared against the baseline from the training corpus

### Selection of corpus for curation and evaluation

To empirically guide curation of entities and synonyms based on the frequency of their appearance in literature we selected a corpus of articles focused on the proteins, protein families, complexes, and molecular events relevant to pathway biocuration (Figure 1B). Specifically, we combined the 3,752 signaling proteins in Reactome [22] with the members of protein families and complexes defined in OpenBEL resource files [23]. From this gene list a corpus of 269,489 papers was assembled by retrieving papers curated for each gene from the Entrez Gene database [24]. Abstracts were obtained from MEDLINE and full texts were downloaded either from the Pubmed Central Open Access subset (in XML or text format), the Pubmed Central Author Manuscript Collection, or via the Elsevier text and data mining API (Table 1).

**Table 1.** Composition of article corpus by source.

| Text Source Type | Number | % |
| --- | --- | --- |
| MEDLINE Abstract | 187,176 | 69.5 |
| Elsevier XML | 35,327 | 13.1 |
| Pubmed Central Open Access Subset XML | 32,113 | 11.9 |
| Pubmed Central Author Manuscript XML | 13,777 | 5.1 |
| No content retrieved | 267 | 0.1 |
| Pubmed Central Open Access Subset text file | 52 | 0.02 |
| **Total** | **269,489** | **100** |

### Event extraction from text using REACH and INDRA

The corpus of ∼270,000 papers was processed with the REACH event extraction software [21], yielding a set of sentences, named entities, and extracted relations (Figure 1B). REACH is built on widely-used methods for syntactic parsing and named-entity recognition: it uses the Stanford CoreNLP parser [25] for syntactic parsing and draws information on biology-specific named entities from Uniprot, InterPro, PFAM, HMDB, ChEBI, Gene Ontology, MeSH, Cellosaurus, ATCC, and CellOntology. As a final step we used the INDRA software [26] to convert events extracted by REACH into INDRA Statements, a format suitable for analyzing and assembling sets of mechanisms into networks and executable models of various kinds.

### Characterizing patterns of grounding errors

The set of entities and events extracted by REACH was used to characterize patterns of grounding errors and prioritize entities and their lexical synonyms for subsequent curation (Figure 1B). Prior to curation, the corpus of articles was divided into two sets: a “training” set and a “test” set consisting of 80% (215,360) and 20% (53,840) of the articles, respectively. The “training” set of articles was processed with REACH in the absence of FamPlex to evaluate baseline grounding accuracy and guide curation. Following curation, the “test” set of articles was processed with a version of REACH incorporating FamPlex. The partitioning of articles was performed to ensure that estimates of grounding accuracy would not be biased toward the specific set of articles used for curation.

### Definition of protein families and complexes and their constituents

Identifiers for protein families and complexes in FamPlex were created by drawing on two resources: 1) identifiers created *de novo* in FamPlex to correspond to named entities encountered in event extraction, and 2) identifiers drawn from the OpenBEL resource. In the first case, identifiers were prioritized by their frequency of occurrence among extracted events, with common entities such as “NF-kappaB”, “Ras”, “PI3-kinase”, “Akt”, etc., accounting for a significant fraction of grounding errors. In the case of OpenBEL, identifiers for protein families and complexes were drawn from the resource files `protein-families.xbel` and `named-complexes.xbel`, accessible via the OpenBEL GitHub repository at https://github.com/OpenBEL/openbel-framework-resources. The full list of all FamPlex identifiers is contained in the text file `entities.csv`.

Members of protein families and complexes are enumerated in the file `relations.csv` using two types of relations: *isa* and *partof*, denoting membership in a family or a complex, respectively (Figure 1A). These relationships can be applied hierarchically to describe multi-level protein subfamily relationships or protein complexes that are hetero-oligomers of subunits belonging to distinct families (Figure 2A). For example, 5’ AMP-activated protein kinase, or AMPK, is a heterotrimeric protein consisting of alpha, beta, and gamma subunits: the alpha and beta subunits comprise families with two isoforms each, and the gamma subunit family has three isoforms. This hierarchical structure can be represented in FamPlex by using a combination of *isa* and *partof* relationships to link the identifiers for the subunit genes to FamPlex-specific identifiers for the subunit families and the full complex (Figure 2A).

**Figure 2.**
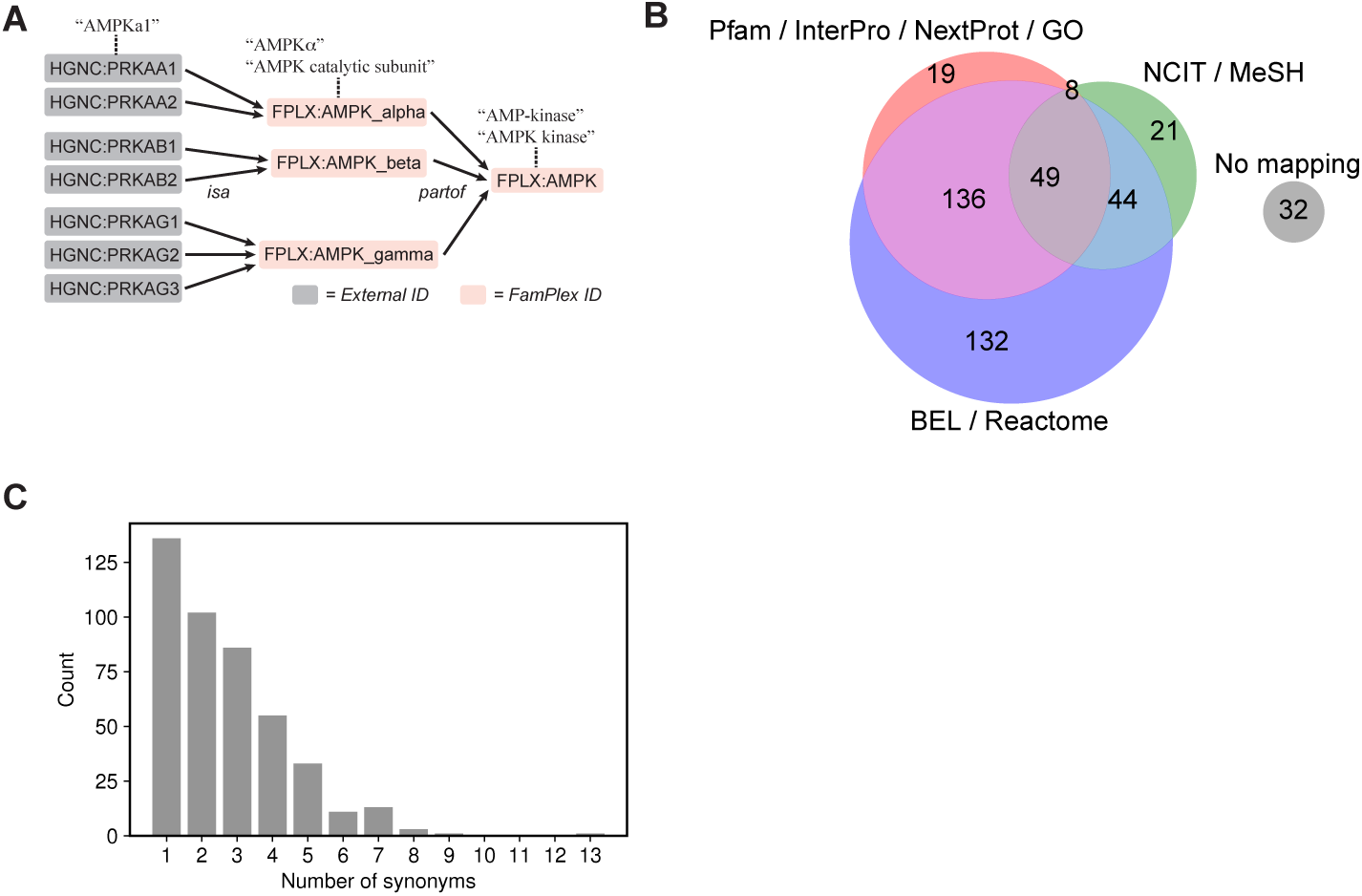
FamPlex links identifiers for families and complexes to members, other databases, and lexical synonyms. **(A)** FamPlex uses *isa* and *partof* predicates to represent the hierarchical relationships between specific genes, families and complexes. Lexical synonyms can be associated with entities at each level. **(B)** Mappings of FamPlex identifiers to outside databases. **(C)** Number of lexical synonyms curated for FamPlex identifiers in the grounding map.

Information on protein family and complex membership was drawn from OpenBEL resource files, HGNC, Reactome, InterPro, and Wikipedia, and manually curated for consistency. Where there were discrepancies among sources about family or complex membership we prioritized what we judged to be the most common usage. For example, the InterPro entry corresponding to the Ras protein family (`IPR020849`) lists 145 human proteins as members, whereas usage in literature and interaction databases recognizes only KRAS, NRAS, and HRAS as family members.

### Mapping FamPlex identifiers to related resources

Entities defined in FamPlex are cross-referenced to corresponding identifiers in other databases and ontologies in the equivalences file (`equivalences.csv`; Figure 1A). Figure 2B shows the subsets of FamPlex identifiers containing mappings to different types of external databases: databases of interactions curated from literature (OpenBEL, Reactome), databases containing specific information about protein families and complexes (PFAM, InterPro, NextProt, and Gene Ontology), and general-purpose biomedical vocabularies (NCIT, MeSH). There are 32 unmapped entries for which no equivalent entry was found in external databases; these entries are implicitly defined in FamPlex by the specific genes that they contain as members.

Identifier mappings between FamPlex and Reactome and InterPro were obtained in a semi-automated fashion. The gene-level members of each FamPlex family and complex were used to query Reactome and InterPro for families and complexes containing these genes. Reactome and InterPro families with equivalent sets of members were incorporated into `equivalences.csv`. Python scripts for generating and updating these mappings are available in the FamPlex GitHub repository at `import/reactome mappings.py` and `import/interpro mappings.py`. Additional identifier mappings to PFAM, NCIT, NextProt, GO and MeSH were collected by entering FamPlex identifiers and lexicalizations into the TRIPS/DRUM web service available at http://trips.ihmc.us/parser/cgi/drum [27]. The TRIPS/DRUM web service returned identifier mappings and their scores based on partial string matches to a variety of databases, which were then manually curated for inclusion in FamPlex.

### Curation of lexical synonyms for entities

Entities defined in FamPlex are associated with lexical synonyms in the grounding map (`grounding map.csv`; Figure 1A). These synonyms allow natural language processing tools to match named entities extracted from text to the protein families and complexes contained in the FamPlex hierarchy.

Lexical synonyms were curated in two ways. First, named entities extracted from the “training” articles read by REACH were sorted by frequency, and named entities corresponding to FamPlex families and complexes were added to the grounding map. Entries were also added to the grounding map for frequently occurring but incorrectly grounded named entities of other types (e.g., proteins, chemicals, and biological processes). For less-frequently encountered families and complexes, synonyms were curated using a different approach: names and synonyms for the gene-level members of families and complexes were used to search the named entities extracted by REACH. Potential matches were identified by fuzzy string matching (Levenshtein distance [28]) using the Python `fuzzywuzzy` package and subsequently manually curated.

Of the 2,076 entries in the FamPlex grounding map, 1,186 map to FamPlex identifiers; the remaining 890 map to frequently occurring proteins, chemicals, and biological processes. The distribution of lexical synonyms across the set of FamPlex identifiers is shown in Figure 2C. The frequently-occurring entities `NFkappaB` and `ERK` have the most synonyms, with 13 and 9, respectively; many other less-frequently occurring entities have only a single synonym. Examples of synonyms for `NFkappaB` include “NF-kB”, “NFkappaB”, and “NF-kappaB TFs”; synonyms for `ERK` include “ERK 1/2”, “ERKs”, and “Extracellular Signal Regulated Kinase”.

### Curation of gene/protein affixes

References to genes and proteins in the literature are often modified by affixes that describe modifications or other context. For example, “mmu-AKT1” and “pAKT1” refer to murine and phosphorylated AKT1, respectively. A list of 137 case-sensitive affixes was tabulated by alphabetically sorting a list of ∼80,000 named entities resulting from event extraction and manually identifying common affix patterns. These affixes were subsequently grouped into six semantic categories (Table 2). The largest category, “experimental context”, contains affixes used to identify the precise variant of a gene used in an experiment; these often refer to protein tags or gene delivery methods. Two of the six categories affect event extraction as well as grounding: “protein state” affixes contain information on modification, location and mutation states, while “inhibition” affixes invert the apparent polarity of an extracted event. For example, a positive regulation event mediated by “BRAF siRNA” actually represents a negative regulation by BRAF itself. The full list of affixes can be found in the CSV file `gene prefixes.csv` (Figure 1A).

**Table 2.** Gene/protein affix types.

| Category | # of affixes | Example |
| --- | --- | --- |
| Experimental context | 63 | eGFP-{Gene name} |
| Protein state | 30 | phospho-{Gene name} |
| Inhibition | 22 | shRNA-{Gene name} |
| Generic descriptor | 12 | proto-oncogene {Gene name} |
| Species | 9 | mmu-{Gene name} |
| mRNA grounding | 1 | {Gene name} mRNA |

### Resource structure and scope

FamPlex comprises 441 families and complexes that together cover 2,040 specific genes through *isa* and *partof* relations. Most FamPlex entries (315) are at the top level of the hierarchy, having no parent entities; 111 entries are at an intermediate level, having both parent and child entities; 15 entities function as placeholders with no parent or child relations currently specified. This latter category consists primarily of functional categories with many potential protein members, e.g., `GTPase`, `Phosphatase`, `Protease`, etc.

The top-level entries vary in terms of the depth of the hierarchy they subsume with the majority of entries (275 in total, two examples being `RAS` and `RAF`) directly being resolved to a set of specific constituent genes. 37 entries have two subsumed levels (for instance `PLC` which subsumes the subfamilies `PLCD`, `PLCG`, and `PLCB`, which in turn subsume a total of nine constituent genes), and 3 entries (`G protein`, `HSP90` and `PI3K`) subsume three levels.

FamPlex entries vary in terms of the number of children they subsume with an average of 6.0 *±* 7.1 children, the large standard deviation indicating the longtailed nature of the distribution. While the median FamPlex entry has 3 children, several entries have a much larger number, including `RAB` (68 children), `Histone` (60 children) and `Cyclin` (31 children).

To characterize the scope and relevance of the different identifiers we quantified the prevalence of each FamPlex entry in PubMed-indexed articles. We conducted PubMed searches for each lexicalization of a given FamPlex entry (using the relatively restrictive “text word” search mode of PubMed to avoid partial matches and matches to meta-information) and counted the total number of unique articles found for each FamPlex entry itself and also for each entry and all its children. The total number of PubMed-indexed articles mentioning one or more FamPlex entries (or children) was 4,012,468, or roughly 14% of all PubMed-indexed literature. The mean number of citations per FamPlex entry was 13,091 *±* 26,733 with a median of 3,034, reflecting a distribution skewed toward a small number of highly cited entries. When including the children of each entry, the number of citations per entry was higher, with a mean of 16,136 *±* 29,491 and a median of 4,929. The most commonly appearing FamPlex entry was `Interferon` with 204,228 associated articles; only 11 FamPlex entries had fewer than 100 associated PubMed citations. Thus, FamPlex covers entities that are frequently mentioned in the biomedical literature.

## Utility and Discussion

### Protein families and complexes appear frequently in events extracted from literature and are often incorrectly grounded

To evaluate baseline grounding performance without FamPlex we manually scored a random sample of 300 named entities generated by running REACH on the training corpus. Entities were categorized by type (protein/gene, family/complex, small molecule, biological process, microRNA, and other/unknown) and the database mappings identified by REACH were scored for correctness (Table 3). Where the entity text alone was insufficient to evaluate grounding accuracy, the sentence in which the entity was embedded was examined in the context of the original paper. We found that references to protein families and complexes were second only to genes and proteins in the frequency of their occurrence in events extracted from text, accounting for 17.7% of all extracted entities (Table 3). Grounding accuracy was substantially lower for families and complexes relative to genes and proteins, with only 15.1% of families and complexes correctly grounded compared to 78.7% for individual proteins (Table 3). The 15% rate of correct grounding for families and complexes reflected accurate matches to identifiers in InterPro or PFAM. Notably, seven of the top ten most frequently occurring *ungrounded* entity texts in the training corpus represented families or complexes (“NF-kappaB”, “ERK1/2”, “mTORC1”, “NFkappaB”, “PDGF”, “IKK”, and “histone H3”; Table 4). Overall, REACH identified a total of 163,428 *unique* named entity strings involved in events, out of which 2,873 were grounded (correctly or incorrectly) to a protein family or complex (1.8%).

**Table 3.**
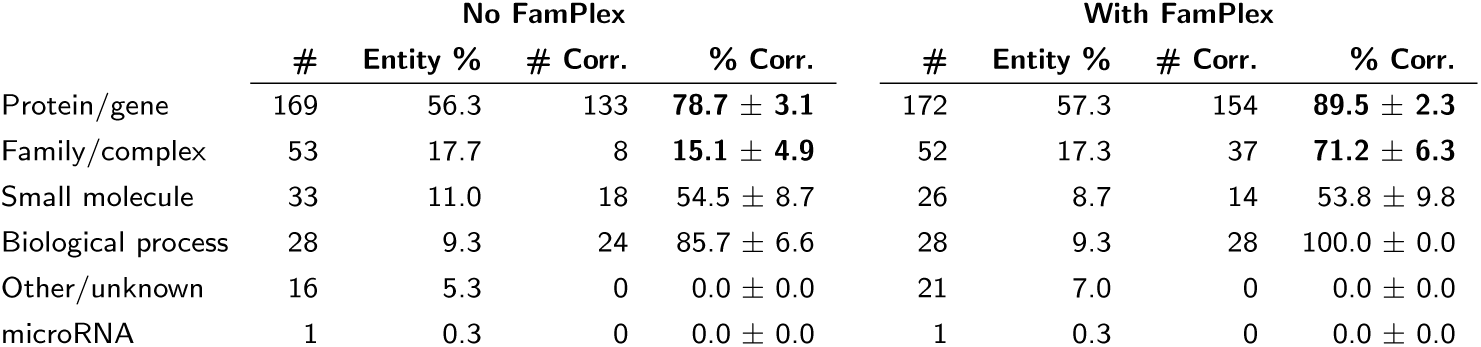
Entity frequency and grounding accuracy for 300 entities, with and without FamPlex. Standard error was calculated using the formula 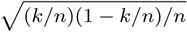 where *k* is the number of samples in the given category and *n* is the total number of samples.

**Table 4.** Top 10 most frequently occurring ungrounded entity texts with and without FamPlex in the training and test corpora, respectively.

| Rank | No FamPlex |  |  | With FamPlex |  |  |
| --- | --- | --- | --- | --- | --- | --- |
|  | Entity Text | Count | % of Total | Entity Text | Count | % of Total |
| 1 | NF-kappaB | 18,381 | 3.78 | PKCzeta | 222 | 0.24 |
| 2 | ERK1/2 | 6,137 | 1.26 | RANTES | 176 | 0.19 |
| 3 | mTORC1 | 2,753 | 0.57 | DC | 169 | 0.18 |
| 4 | NFkappaB | 2,425 | 0.50 | LDL | 168 | 0.18 |
| 5 | c-Jun | 2,369 | 0.49 | IgE | 152 | 0.17 |
| 6 | antigen | 1,724 | 0.35 | SDF-1-alpha | 141 | 0.15 |
| 7 | PDGF | 1,626 | 0.33 | receptor | 128 | 0.14 |
| 8 | IKK | 1,542 | 0.32 | beta1 integrin | 127 | 0.14 |
| 9 | c-Src | 1,362 | 0.28 | p38alpha | 126 | 0.14 |
| 10 | histone H3 | 1,347 | 0.28 | CD4+ | 124 | 0.14 |

Close inspection of errors made by REACH in grounding frequently-occurring families and complexes in the absence of FamPlex revealed the causes of both missing and incorrect groundings. Missing groundings occurred when named entities corresponding to families and complexes had no associated identifiers, indicating a failure to find a match in any database. This was true of the entity “Ras”, as well as the most frequently occurring family-level entity, “NF-kappaB”.

On the other hand, incorrect grounding of family-level entities occurred due to exact (but spurious) matches to obscure synonyms for other genes listed in Uniprot or HGNC. In some cases these genes were unrelated to the family but had synonyms shadowing the family name: for example, “ERK” and “Cyclin” were grounded to the human genes *EPHB2* (Uniprot `P29323`) and *PCNA* (Uniprot `P12004`) due to the presence of these strings as synonyms. Another class of grounding errors involved the matching of a string representing the basename of a human protein family to the single ortholog of the family in different organism. Representative examples include the misgrounding of “AKT” to the *Dictyostelium discoideum* gene *pkbA* and of “JNK” to the *Drosophila melanogaster* gene *bsk*, both of these listing the human gene family name as synonyms.

The most common ungrounded strings (those in the highest percentile by frequency of occurrence) accounted for a surprisingly large proportion of the *overall* number of ungrounded string occurrences, as shown by the orange curve in Figure 3A. The deviation of this curve from a uniform distribution (shown by the dotted gray line in Figure 3A) arises because the empirical distribution of ungrounded entities is highly skewed, with a small number of very common entities accounting for a large percentage of occurrences. For example, half of all ungrounded string occurrences in the training corpus involved the top 2.4% most frequently occurring strings (2,666 distinct strings). This explains why curation that is focused specifically on frequently occurring misgrounded entities has the potential to substantially improve overall grounding and reading performance.

**Figure 3.**
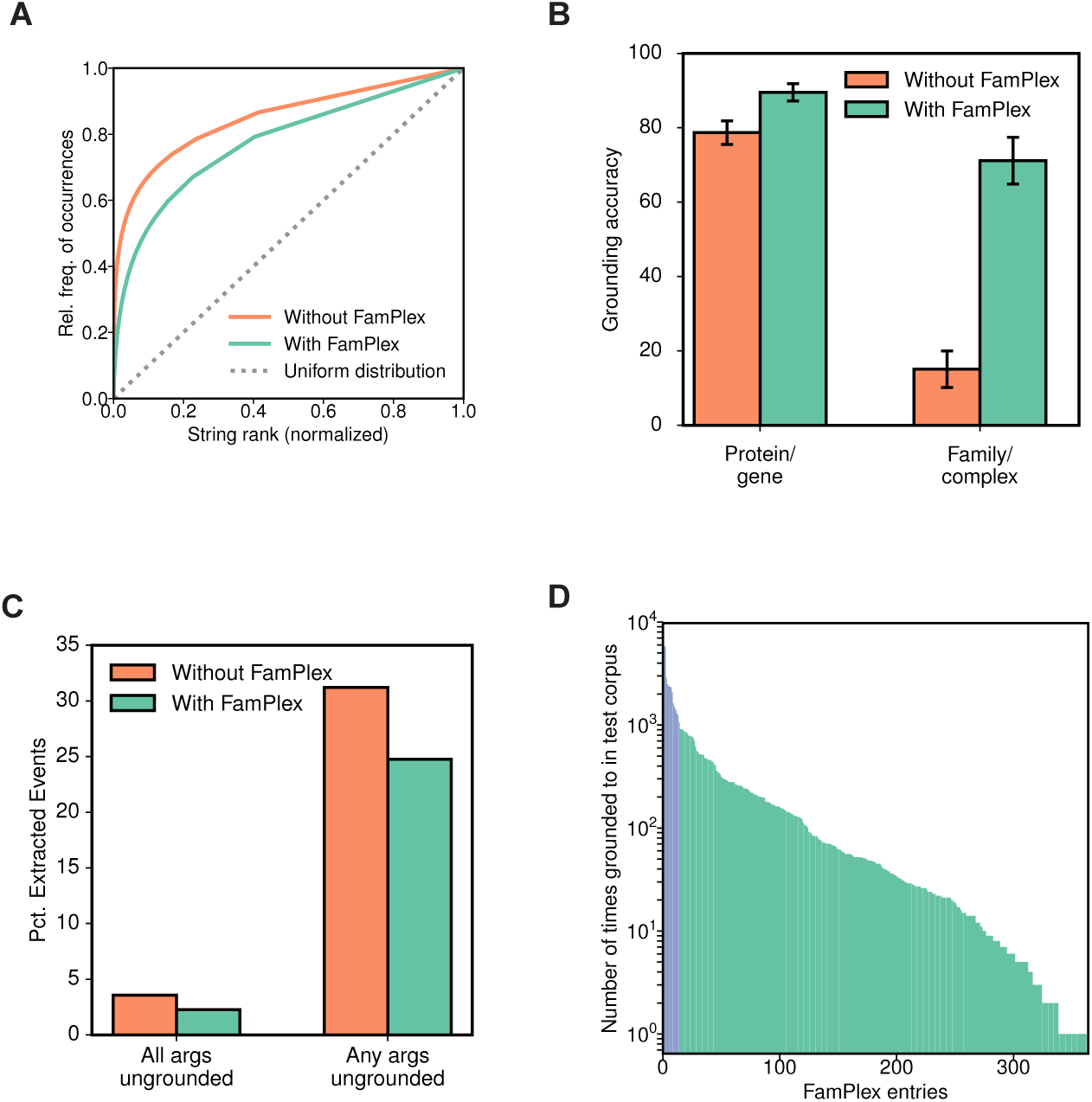
FamPlex improves grounding accuracy. **(A)** Cumulative occurrences of ungrounded entities by frequency of the entity text. Deviation from the dotted gray line, representing a uniform frequency distribution, indicates the extent to which a small number of frequently occurring entities account for a disproportionate share of missed groundings. **(B)** Improvements in grounding accuracy for proteins/genes and families/complexes, with and without the use of FamPlex. **(C)** Reduction in the proportion of extracted events containing ungrounded entities, with and without FamPlex. **(D)** Number of groundings to FamPlex identifiers in the test corpus. The 15 most frequent identifiers account for 50% of all groundings and are shown in blue.

### Use of FamPlex in text mining improves grounding and relationship resolution for protein families and complexes in two event extraction systems

Following the manual curation of FamPlex identifiers and associated synonyms and the integration of FamPlex into REACH and INDRA, we performed a second evaluation on a random sample of 300 named entities drawn from the results of processing the test corpus (Table 3). The frequency of entity types was comparable between the training and test samples, with proteins/genes and families/complexes accounting for roughly three-quarters of all entities. Improvements in grounding were substantial for both classes, with grounding accuracy for families and complexes rising from 15% to 71% (Figure 3B; Table 3). Grounding accuracy for proteins and genes increased from 79% to 90%, an improvement attributable to the curation of synonyms for frequently occurring proteins. With the incorporation of FamPlex, the overall percentage of unique entity strings grounded to protein family or complex identifiers doubled relative to the training corpus, with REACH grounding 2,080 of 57,088 unique entities to a FamPlex, InterPro or PFAM identifier (3.6%).

An analysis of the distribution of the remaining ungrounded entities showed that FamPlex addressed a substantial proportion of the most frequently occurring grounding failures (Figure 3A, green curve). As shown in Table 4, the top ten most frequently occurring ungrounded entities in the test set occur at a lower overall frequency and include a functional category (“receptor”) but no specific protein families or complexes. To examine the impact of grounding improvements at the level of extracted events, we calculated the proportion of events consisting either of any or all ungrounded entities, and found that both metrics improved with the use of FamPlex (Figure 3C). These measures, which deal only with event entities that were *ungrounded*, represent an underestimate of the overall improvement in grounding because they do not account for cases in which entities were grounded to the *wrong* identifier in the absence of FamPlex.

To characterize whether improvements in grounding were driven by a small subset of frequently-occurring entities in FamPlex or were more broadly distributed across families and complexes, we counted the occurrences of mappings to each FamPlex identifier in events extracted from the test corpus. We found that the 15 most frequently-referenced FamPlex identifiers accounted for 50% of all FamPlex groundings (blue bars in Figure 3D); the top five are shown in Table 5. At the same time, 363 of the 441 FamPlex identifiers were mapped to text at least once, suggesting that the great majority of identifiers and lexical synonyms in FamPlex are useful for improving grounding (Figure 3D).

**Table 5.**
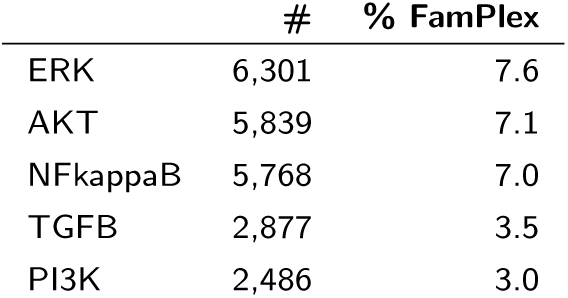
FamPlex entries most frequently grounded to in test corpus, with the absolute number of times grounded to in the test corpus and the percentage normalized to all FamPlex groundings.

As a second means to evaluate FamPlex we used the TRIPS/DRUM reading system [27]. Unlike REACH, which uses strict string matching against a set of dictionaries, TRIPS uses soft matching to provide a ranked, scored list of groundings for each named entity. Relevant dictionaries used by TRIPS include PFAM and NextProt for protein families, GO for protein complexes and NCIT for both.

We compiled two versions of TRIPS, one in which FamPlex was included as a grounding resource, and one in which it was omitted. Since the throughput of TRIPS is substantially lower than that of REACH, we selected a random sample of 100 abstracts from the combined training and test set for reading with and without FamPlex. We then manually curated 500 randomly sampled entities appearing in TRIPS extractions, determining whether each entity represented a protein family or complex, and if so, whether: (i) the *top scoring* grounding match was correct, and (ii) *any* of the grounding matches were correct. In contrast to our evaluation of entity grounding in REACH, in which the curated entities were limited to arguments of events, here we considered *all* entities identified in text by TRIPS as candidate families or complexes for curation. This broader pool of candidate entities included names of cell lines, organisms, biological processes, etc., and therefore also a smaller proportion of molecular entities such as families and complexes.

In the case of TRIPS without FamPlex, 36 of 500 entities sampled from the TRIPS output corresponded to families or complexes. Of these, we found that the top scoring grounding was correct for 23 (64%); 29 entities (81%) had at least one correct grounding. The higher baseline accuracy of family/complex grounding in comparison with REACH likely reflects broader coverage of relevant identifiers due to the inclusion of NextProt and NCIT (used by TRIPS but not by REACH) and the more robust but computationally costly soft-matching and ranking procedure used for grounding. While no single resource accounted for the majority of all matches, top-scoring matches were roughly equally distributed between NCIT and NextProt. Moreover, of the 17 entities that were correctly grounded in NCIT, 7 (41%) had no identified child concepts, making it impossible to link these families and complexes to constituent genes. Thus, while TRIPS was more successful than REACH at finding relevant groundings for families and complexes in the absence of FamPlex, the multiplicity of alternative groundings and the unresolved nature of these terms in the ontologies used posed a distinct problem, that of relationship resolution.

Incorporating FamPlex into TRIPS improved both the accuracy and consistency of grounding. In a sample of 500 entities extracted by TRIPS using FamPlex, 33 corresponded to families and complexes; the top-scoring grounding was correct for 26 (79%) of these and a further four (91% overall) had at least one correct grounding. While the small sample sizes limit quantitative conclusions about the degree of improvement, we noted that in 18 of 26 (69%) cases in which the top-scoring grounding was correct, it was grounded to a FamPlex identifier, and in 20 of 26 (77%) a FamPlex grounding was among the top two matches. This indicates that FamPlex identifiers and lexicalizations have a higher coverage for families and complexes encountered in text by TRIPS than other resources used, allowing for more consistent relationship resolution and integration of information.

### FamPlex includes a large majority of families and complexes annotated by human curators in text

In addition to the evaluations of grounding *precision* described above, we sought to establish a measure of the *recall* of FamPlex in terms of its coverage of relevant families and complexes in a manually curated dataset. Evaluations solely against machine reading output, as described above, do not provide a true recall measure because the readers extract only a subset of the events and entities from the underlying text.

To evaluate recall we used the dataset prepared for the bioentity normalization task from Biocreative VI Task 1.1 (http://www.biocreative.org/tasks/biocreative-vi/track-1/). The dataset, drawn from the EMBO SourceData annotation project [29], contains a corpus of entity text strings from figure legends in published papers, most of which have been annotated with database identifiers by human curators. Our aim was to evaluate the extent to which FamPlex incorporates identifiers and lexicalizations for the family and complex-level entities identified in text by human curators.

Inspection of the Biocreative dataset revealed that curators annotated family- and complex-level strings in multiple ways: to a single gene, multiple genes, or simply left ungrounded. We therefore partitioned the annotation data into multiple subsets for the purposes of evaluation (Table 6). The first of these was the subset of 19,228 entities grounded to human Uniprot or NCBI gene identifiers, which we denote Annotation Subset 1 (AS1; 18.7% of the total). Of these, 2,439 entities (2.4% overall) were grounded to *multiple* human gene or protein identifiers; these therefore correspond to gene families or protein complexes (denoted AS2). We also drew from “ungrounded” entities, i.e., annotations labeled “gene” or “protein” but lacking identifiers. A large majority of these represented experimental elements or protein tags, e.g. “GFP”, “FLAG”, “GST”, etc. To streamline curation, we filtered ungrounded entities against the affixes included in FamPlex; a high proportion of ungrounded entities (8,250 of 14,227, or 58%) had matches in the FamPlex affixes list in `gene prefixes.csv`, leaving 5,977 entities for further curation, a subset denoted AS3 (Table 6).

**Table 6.** Subsets of the Biocreative VI entity normalization dataset relevant to the FamPlex evaluation. Entities evaluated against FamPlex were drawn from the categories highlighted in bold.

| Annotation Category | # | % of total |
| --- | --- | --- |
| All annotations | 102,717 | 100.0 |
| Grounded to gene/protein | 44,576 | 43.4 |
| **Grounded to human gene/protein (AS1)** | 19,228 | 18.7 |
| **Grounded to multiple human genes/proteins (AS2)** | 2,439 | 2.4 |
| Ungrounded gene/protein | 14,227 | 13.9 |
| Ungrounded gene/protein matching FamPlex affix | 8,250 | 8.0 |
| **Ungrounded gene/protein not matching FamPlex affix (AS3)** | 5,977 | 5.8 |

An initial round of scoring focused exclusively on identifying the proportion of the 2,439 entities in AS2 (the subset containing multiple gene/protein groundings) covered by FamPlex; we found that 1,908 (78%) had case-insensitive matches in the FamPlex grounding map. Of the remaining 531 unmatched entities (representing 109 unique strings), manual curation indicated that 51 corresponded to noncoding RNAs and were excluded, leaving 2,388 entities (1,908 + 480) with multiple gene/protein groundings. Of the remaining 480 entities representing proteins, manual curation indicated that 97 had corresponding identifiers in FamPlex. We therefore calculated that FamPlex contained both string matches *and* identifiers for 79.9% of the entity texts in AS2, and identifiers but not string matches for a slightly higher proportion (84%; Table 7).

**Table 7.** Extent of FamPlex family/complex coverage evaluated against subsets of the Biocreative VI entity normalization dataset.

| Annotations Scored | String Matches | Corresponding IDs |
| --- | --- | --- |
| Multiple gene/protein groundings (AS2) | 1908 / 2388 (79.9%) | 2005 / 2388 (84.0%) |
| Families curated from random sample of AS1 + AS3 | 89 / 109 (81.7%) | 96 / 109 (88.1%) |

Because families were not always grounded to multiple gene/protein identifiers by human curators, we performed a second evaluation in which we manually curated a random sample of entities drawn from AS1 + AS3. Of 764 curated entity strings, 109 were found to be synonyms for protein families or complexes (note that, unlike in the evaluation against AS2 above, this assessment was made independently of the annotations contained in the dataset). As in the previous evaluation, these were scored for the presence of string matches and/or corresponding IDs in FamPlex, yielding similar figures of 81.7% and 88.1%, respectively (Table 7). Taken together, these results demonstrate that FamPlex incorporates identifiers and lexical synonyms for a large proportion of the families and complexes relevant to manual biocuration tasks from literature.

### FamPlex resolves hierarchical relationships in extracted events

A key feature of FamPlex is that it allows for relationship resolution not only “horizontally” (between different databases) but also “vertically” (between genes, families, complexes, and any intermediate sets involving these elements). Lexical synonyms can be defined at all levels in the FamPlex hierarchy (Figure 2A). The combination of a hierarchical representation with a mapping of entities to text at each level allows information about biological interactions to be correctly organized and cross-referenced.

For example, the FamPlex family `PLC`, representing the family of phospholipase C enzymes, contains both individual genes (e.g., *PLCE1*) and FamPlex subfamilies (e.g., `PLCG`, a sub-family consisting of the genes *PLCG1* and *PLCG2*) as members (Figure 4A). In results from the test corpus we found descriptions of meaningful biochemical mechanisms associated with all three levels of this hierarchy—family, subfamily, and genes (Figure 4A). Moreover, relevant events were extracted for 12 of the 15 entities in the phospholipase C entity hierarchy, demonstrating the diversity of available mechanistic information and the importance of relationship resolution.

**Figure 4.**
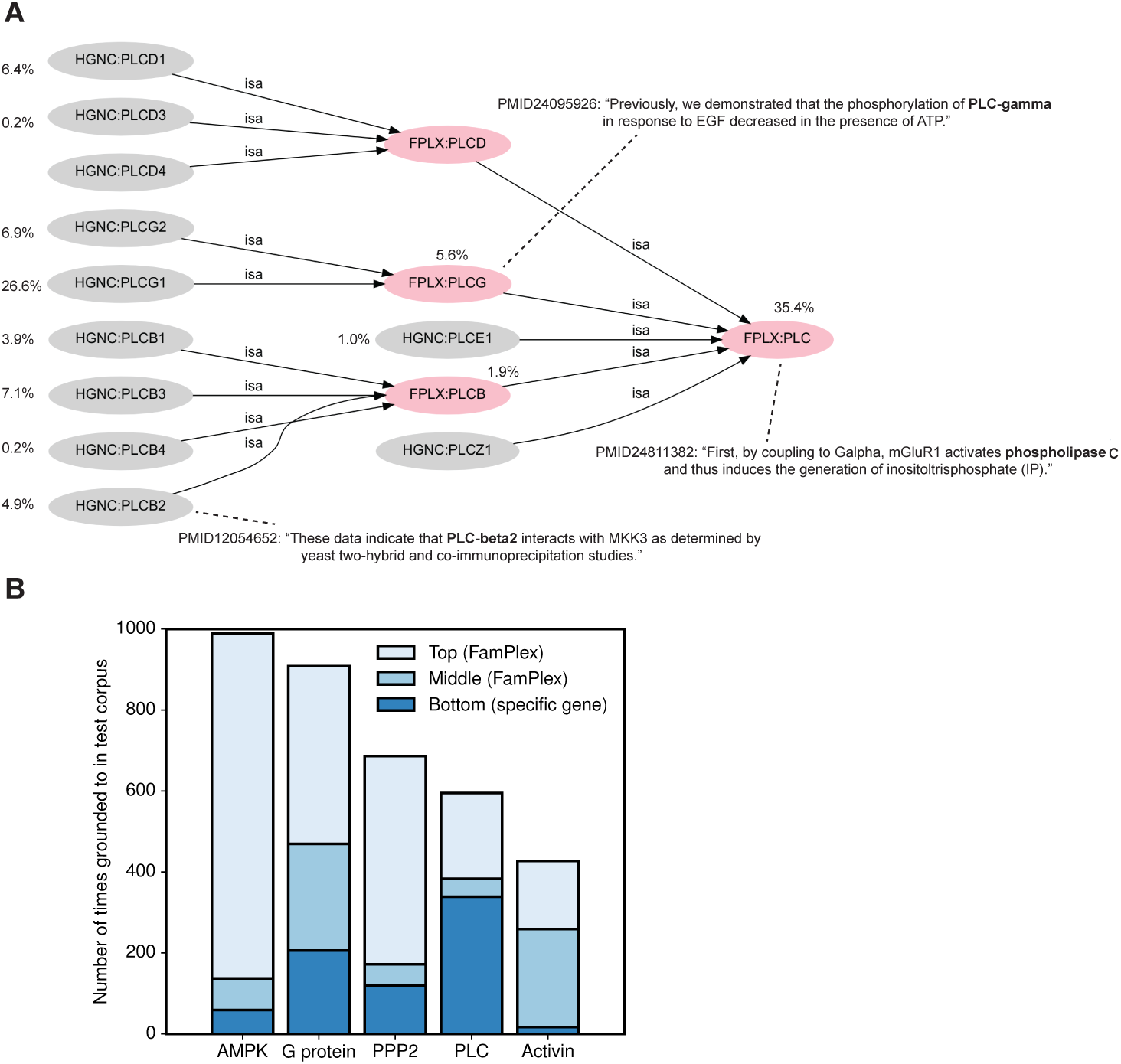
FamPlex facilitates hierarchical resolution of extracted information. **(A)** Hierarchical organization of the phospholipase C protein family (FamPlex identifier `PLC`) along with the proportion of occurrences of each member in the test corpus and examples of sentences yielding information at the different levels. Pink nodes indicate FamPlex families; gray nodes indicate genes. **(B)** Proportion of groundings in the test corpus to gene-level, intermediate-level, or top-level entities for five multi-level families/complexes in FamPlex.

To characterize the relevance of multi-level relationship resolution more broadly, we counted the number of times a named entity identified by REACH in the test corpus was mapped to a FamPlex identifier at three or more hierarchical levels: the gene level (lowest), the top-level family or complex (highest), and any intermediate level. Distributions of groundings for five FamPlex entries with three or more entity levels are shown in Figure 4B. Overall, we found that 33 top-level FamPlex entries (i.e. ones that are not subsumed through an *isa* or *partof* relation by another FamPlex entry) were associated with groundings at three or more distinct levels, and 242 top-level FamPlex entries had groundings at two levels (i.e. grounding to the FamPlex entry itself and its constituent genes), showing that gene functions are commonly discussed across multiple levels of specificity.

We also found that the identifier level used most frequently for grounding differed among protein families and complexes, limiting generalizations about the relative priority of gene-vs. family-level grounding for event extraction. For example, for `AMPK`, the majority of references in the literature were to the top-level AMPK complex, with a relatively small fraction of references to constituent genes or intermediates. On the other hand, most mappings to the family representing Phospholipase C (`PLC` in FamPlex) were to constituent genes such as *PLCG1, PLCD1*, etc. Finally, for the family of Activins (hetero- and homo-dimers of the transforming growth factor beta family, `Activin` in FamPlex), most references were to specific dimer subtypes—Activin A, Activin AB and Activin B—which are found at an intermediate level in the FamPlex hierarchy.

### Comparison of FamPlex with other resources

FamPlex bears similarities to three types of existing resources. The first of these are large, systematic assemblies of protein families derived from sequence and domain analysis; this set includes PFAM, InterPro, and Homologene. As a curated resource, FamPlex is less comprehensive, since it includes only human genes and focuses primarily on gene families and lexicalizations that are described in existing literature. However, FamPlex includes complexes as well as families, based on the observation that these high-level groupings of proteins are often interwoven in discussions of gene function (e.g., “AMPK” and “AMPK-alpha”; Figure 2A). FamPlex also provides lexical synonyms for families and complexes, a feature generally absent from large protein family databases.

A second class of comparable resources are the taxonomies of protein families and complexes defined as part of biocuration projects or tools; examples include Reactome, SIGNOR, and OpenBEL. These taxonomies are designed to meet the need of biocurators to specify mechanistic interactions at the family or complex level. Of these resources, we found the families and complexes defined by OpenBEL to be the most systematic and reusable, and we therefore drew heavily on OpenBEL in the construction of FamPlex. FamPlex differs from the families and complexes defined in resources such as Reactome, SIGNOR and OpenBEL in three important ways: (i) it includes an extensive set of lexicalizations to assist in grounding, (ii) it enumerates equivalent family/complex identifiers *between* many of these resources, allowing for mechanistic information to be integrated at the family/complex level, and (iii) it allows for a *multi-level* entity hierarchy corresponding to the terms and concepts used in the literature.

The third category of related resources are biomedical ontologies such as GO and terminology resources such as NCIT and MeSH. While these resources are the most broadly extensive and often contain synonyms for concepts, they have uneven coverage of protein families and complexes specifically. In addition (as described in our evaluation of grounding to NCIT in the TRIPS reading system) many identifiers representing protein families and complexes do not incorporate child concepts at the gene level, limiting their value for relationship resolution.

Thus, while FamPlex draws on and provides cross-references to all three classes of resources described above, it differs from all of them in providing a consistent, multi-level taxonomy of human protein families and complexes that is suitable for grounding and relationship resolution in text mining and biocuration.

### Limitations

The relatively high recall achieved by FamPlex on the Biocreative entity normalization dataset suggests that it provides substantial coverage of relevant protein families, complexes and their lexical synonyms. However, it is not exhaustive. Further empirically-guided curation of the identifiers and grounding map is likely to improve grounding precision and recall still further, and with additional work mappings to other ontologies can be made more comprehensive.

FamPlex does not directly address the problem of *ambiguity*, selecting among multiple alternative groundings for the same entity. For example, “MEK” can refer to the family of MAPK/ERK Kinases or to the solvent methyl ethyl ketone. Resolving such ambiguities requires an examination of the named entity in the broader context of the sentence or article [30]. However, the use of FamPlex does increase the likelihood that relevant groundings to protein families will not be missed, and can therefore be considered alongside alternative groundings during an ambiguity resolution procedure.

### Accessibility and Extensibility

We chose CSV files as the primary format for FamPlex to maximize accessibility and extensibility. CSV files can be opened and edited in any spreadsheet program or text editor, allowing biologists with no background in literature mining to assist in the curation of the grounding map or create mappings to other resources. Because the files are hosted on GitHub, other users can easily fork and make use-case specific extensions or other contributions that can be merged back into the main repository. In addition to the CSV files, FamPlex includes an Open Biomedical Ontologies (OBO) [31] export feature to facilitate integration into OBO-based workflows. Fam-Plex relations and mappings have been integrated into the TRIPS/DRUM reading system [27] via OBO-exported content.

## Conclusions

In this paper we describe the challenge posed by protein families and multi-protein complexes for machine reading of the biomedical literature. We introduce Fam-Plex, a new lexical and ontological resource that addresses these challenges and improves grounding and relationship resolution in two different reading systems [21, 27]. FamPlex fills a gap in existing bioinformatics resources, linking information about families and complexes in protein and pathway databases to a set of lexical synonyms that occur with high frequency in the scientific literature. Empirical evaluation shows that the hierarchical organization of FamPlex enables the integration of mechanistic information about gene families, complexes, and their individual subunits. This is important because information about biochemical mechanisms is often reported in terms of classes of entities whereas large-scale profiling experiments yield data about individual genes and proteins. FamPlex therefore supports the broader goal of making text mining a key contributor to the process of obtaining biological insight from high throughput -omic data by drawing on relevant mechanistic knowledge. We speculate that similar resources for resolving hierarchical relationships among entities could be useful in other domains of machine reading and natural language processing.

## Declarations

### Ethics approval and consent to participate.

Not applicable.

### Consent for publication

Not applicable.

### Availability of data and material

The datasets generated and analyzed during the current study, as well as the source code used to generate results is available in the repository https://github.com/sorgerlab/famplex paper. FamPlex is available at https://github.com/sorgerlab/famplex under the Creative Commons CC0 license.

### Authors’ contributions

JAB and BMG conceived and implemented the resource. JAB, BMG and PKS wrote the paper.

## Acknowledgements

The authors would like to thank Tom Hicks, Mihai Surdeanu and Lucian Galescu for useful discussions and assistance with REACH and TRIPS, and Petar Todorov, Bobby Sheehan, Lily Chylek, Jeremy Muhlich and Isabel Latorre for their contributions to entity curation.

## Abbreviations

API: Application Programming Interface
CSV: Comma-Separated Variable
DRUM: Deep Reader for Understanding Mechanisms
HGNC: HUGO Gene Nomenclature Committee
HMDB: Human Metabolome Database
INDRA: Integrated Network and Dynamical Reasoning Assembler
MeSH: Medical Subject Headings
NCBO: National Center for Biomedical Ontology
NCIT: National Cancer Institute Thesaurus
NER: Named Entity Recognition
OBO: Open Biological and Biomedical Ontology
REACH: Reading and Assembling Contextual and Holistic Mechanisms
SIGNOR: Signaling Network Open Resource
TRIPS: The Rochester Interactive Planning System.

